# Common host variation drives malaria parasite fitness in healthy human red cells

**DOI:** 10.1101/2020.10.08.332494

**Authors:** Emily R Ebel, Frans A Kuypers, Carrie Lin, Dmitri A Petrov, Elizabeth S Egan

## Abstract

The replication of *Plasmodium falciparum* parasites within red blood cells (RBCs) causes severe disease in humans, especially in Africa. The influence of host RBC variation on parasite replication is largely uncharted, aside from a handful of deleterious alleles like sickle cell. Here, we integrated analyses of exome sequencing, RBC phenotyping, and parasite fitness assays on blood from 122 individuals, most with African ancestry. In donors lacking alleles for hemoglobinopathies or G6PD deficiency, RBC phenotypes including size, deformability, and hydration status explained 21-38% of the variation in parasite growth rate. With the addition of non-pathogenic polymorphisms in 14 RBC proteins including *SPTA1, PIEZO1*, and *ATP2B4*, our model explained 73-82% of the variation in parasite growth rate. Interestingly, we observed little evidence for divergent selection on this variation between Africans and Europeans. These findings suggest a model in which widespread, non-pathogenic variation in a moderate number of genes strongly modulates *P. falciparum* fitness in RBCs.

## INTRODUCTION

Malaria caused by the replication of *Plasmodium falciparum* parasites in red blood cells (RBCs) kills hundreds of thousands of children each year (WHO, 2019). In each 48-hour cycle of blood stage malaria, parasites must deform RBC membranes to invade them(Koch *et al*., 2017; Kariuki *et al*., 2020); consume hemoglobin and tolerate the resulting oxidative stress (Francis, Sullivan and Goldberg, 1997); multiply to displace half the RBC volume (Hanssen *et al*., 2012); and remodel the RBC membrane to avoid immune detection (Zhang *et al*., 2015). Consequently, genetic disorders that alter aspects of RBC biology are well known to influence malaria susceptibility (Kwiatkowski, 2005). For example, sickle cell trait impairs parasite growth by altering hemoglobin polymerization at low oxygen tension (Pasvol, Weatherall and Wilson, 1978; Archer *et al*., 2018), while deficiency of the G6PD enzyme involved in oxidative stress tolerance is thought to make parasitized RBCs more susceptible to breakdown (Ruwende and Hill, 1998). Aside from these diseases, however, the genetic basis of malaria susceptibility remains mostly unknown. Several large genome-wide association studies (GWAS) have recently made progress on this question by identifying a few dozen loci that collectively explain up to 11% of the heritability of the risk of severe versus uncomplicated malaria (Timmann *et al*., 2012; Rockett *et al*., 2014; Band *et al*., 2015; Leffler *et al*., 2017; Ndila *et al*., 2018; MGE Network, 2019a). Eleven of these loci are in or near genes expressed predominantly in RBCs, and one new GWAS variant has since been shown to regulate expression of the *ATP2B4* calcium channel (Zámbó *et al*., 2017) and to be associated with RBC dehydration (Li *et al*., 2013). However, most other RBC loci that have been identified by GWAS are not novel, and a functional link between *ATP2B4* and *P. falciparum* replication has yet to be demonstrated. More discoveries about the influence of RBC variation on malaria susceptibility are not expected from similar GWAS (Boyle, Li and Pritchard, 2017; MGE Network, 2019b), in part because severe malaria is a complex phenotype that depends on a combination of factors from RBCs, the vascular endothelium, the immune system, and the environment (Mackinnon *et al*., 2005; De Mendonça, Goncalves and Barral-Netto, 2012). Alternate approaches are therefore needed to discover more genetic variation that impacts the replication of malaria parasites in human RBCs.

RBC phenotypes like mean cell volume (MCV), hemoglobin content (HGB/MCH), and antigenic blood type vary widely within and between human populations (Lo *et al*., 2011; Cooling, 2015; Canela-Xandri, Rawlik and Tenesa, 2018). 40-90% of the variation in common RBC indices is heritable (Whitfield, Martin and Rao, 1985; Evans, Frazer and Martin, 1999; Pilia *et al*., 2006), and GWAS have demonstrated that these phenotypes have polygenic bases (van der Harst *et al*., 2012; Astle *et al*., 2016; Chami *et al*., 2016; Chen *et al*., 2020; Vuckovic *et al*., 2020). Average hemoglobin levels, hematocrit, and especially RBC membrane fragility have also been shown to differ between African and European populations, likely for genetic reasons (Garn, 1981; Perry *et al*., 1992; Beutler and West, 2005; Kanias *et al*., 2017; Page *et al*., 2017). Although African populations are highly genetically diverse(Campbell and Tishkoff, 2008), some of these differences between Africans and Europeans are explained by RBC disease alleles (Beutler and West, 2005; Lo *et al*., 2011; Kanias *et al*., 2017) that have been widely selected across Africa for their protective effects on malaria. It remains untested whether other extensive phenotypic and genetic diversity in RBCs, aside from these disease alleles, also influences malaria susceptibility and has been similarly shaped by malaria selection.

Here, we approach these questions by performing exome sequencing and extensive RBC phenotyping on blood samples from a diverse human cohort of 122 individuals. We show that *P. falciparum* fitness varies widely among donor cells *in vitro*, with the distributions of parasite phenotypes in ‘healthy’ RBCs overlapping those from RBCs carrying classic disease alleles. We apply Lasso variable selection to identify a small set of alleles and phenotypes that strongly predict parasite fitness outside of the context of RBC disease, highlighting RBC membrane and dehydration genes and phenotypes as key to modulating *P. falciparum* fitness. However, we find no evidence that non-pathogenic alleles or phenotypes that confer parasite protection are associated with African ancestry. Overall, these findings advance our understanding of the origin and function of common RBC variation and suggest new targets for therapeutic intervention for malaria.

## RESULTS

### Many healthy blood donors with African ancestry carry alleles for RBC disease

We collected blood samples from 121 donors with no known history of blood disorders, most of whom self-identified as having recent African ancestry (Figure 1A). As a positive control, we also sampled a patient with hereditary elliptocytosis (HE), a polygenic condition characterized by extremely fragile RBC membranes that strongly inhibit *P. falciparum* growth (Schulman *et al*., 1990; Facer, 1995; Dhermy, Schrével and Lecomte, 2007; Gallagher, 2013). We performed whole-exome sequencing, both to check for the presence of known RBC disease alleles and to confirm the population genetic ancestry of our donors. A principal component analysis of more than 35,000 unlinked, exomic SNPs showed that most donors fell along a continuum from African to European ancestry, as defined by data from the 1000 Genomes Project (Figure 1A). We found that 16% of the healthy donors carried pathogenic hemoglobin alleles (Figure 1B), including 5 heterozygotes for hemoglobin S (HbAS), 4 heterozygotes for hemoglobin C (HbAC), and 11 individuals with one or two copies of an *HBA2* deletion causing α-thalassemia (Galanello and Cao, 2011). We also scored eight polymorphisms in *G6PD* that have been functionally associated with various degrees of G6PD deficiency (Yoshida, Beutler and Motulsky, 1971; Clarke *et al*., 2017) and found that 32% of the study population carried at least one, including 12 of the 20 donors with hemoglobinopathies. Among those with wild-type hemoglobin, we identified one individual with polymorphisms associated with severe G6PD deficiency (>60% loss of function) and 23 with polymorphisms associated with mild to medium deficiency (<42% loss of function). To our knowledge, we detected no alleles linked to other monogenic RBC disorders, including β-thalassemia or xerocytosis (Cao and Galanello, 2010; Glogowska *et al*., 2017). We therefore classified the remaining 73 donors as ‘non-carriers’ of known disease alleles for the purposes of this work.

**Figure 1.**
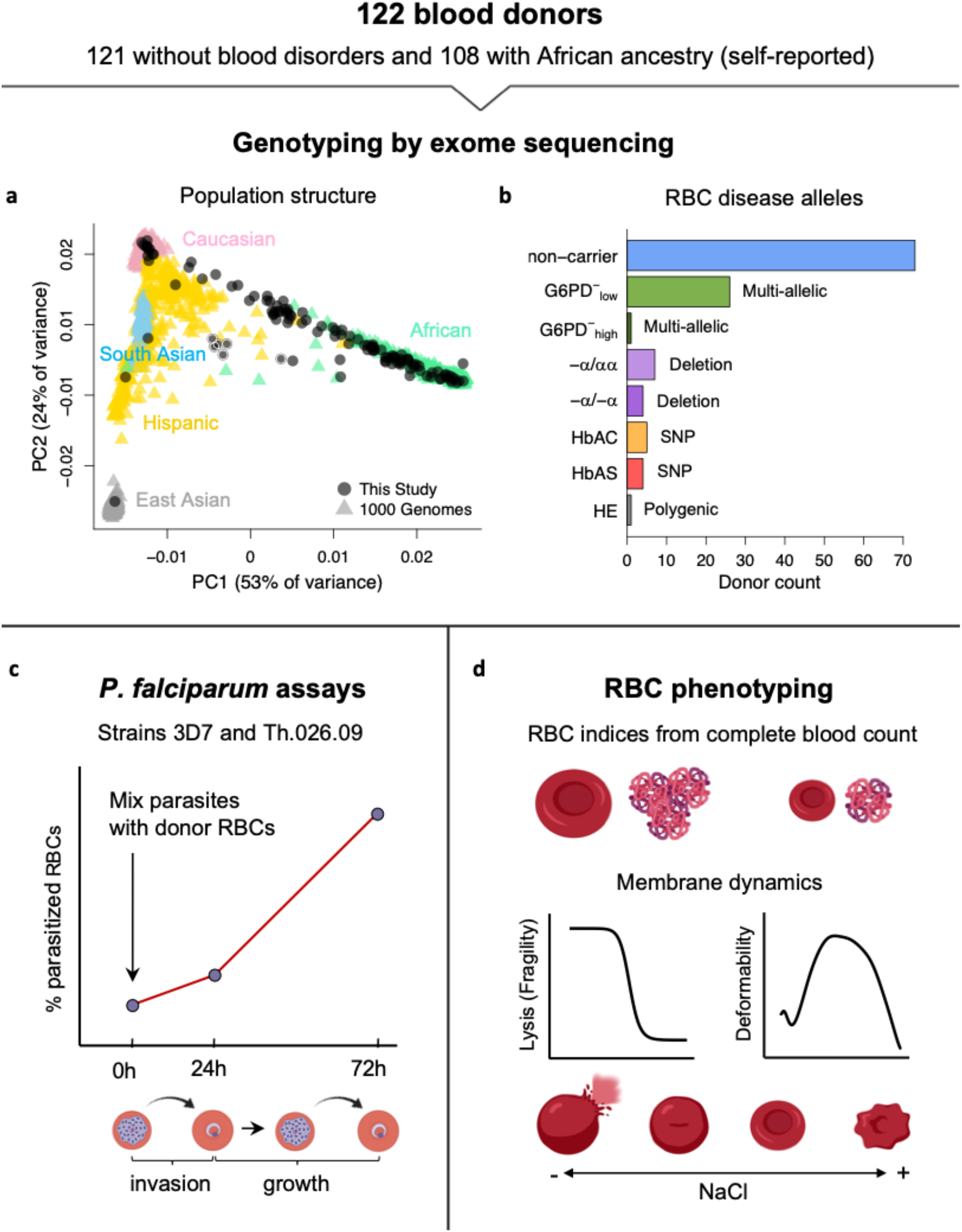
Overview of study design and blood donors. **(a)** PCA of genetic variation across 35,759 unlinked exome SNPs. Samples collected in this study are plotted on coordinate space derived from 1000 Genomes reference populations. Points with white borders represent the six related individuals in this study. **(b)** Exome sequencing revealed that 37% of donors carried alleles for RBC disorders linked to *P. falciparum* resistance. Individuals with >1 disease allele were assigned to the category of the most severe condition. **non-carrier:** Donor without any of the following alleles or conditions. **G6PD-_low_:** Mild to medium G6PD deficiency (<42% loss of function). **G6PD-_high:_** Severe G6PD deficiency (>60% loss of function). **-α/αα:** heterozygous HBA2 deletion, or alpha thalassemia minima. **-α/-α:** homozygous HBA2 deletion, or alpha thalassemia trait. **HbAC**: heterozygous HBB:E7K, or hemoglobin C trait. **HbAS:** heterozygous HBB:E7V, or sickle cell trait. **HE:** hereditary elliptocytosis. **(c)** Parasite replication rate in donor RBCs was measured via flow cytometry at three timepoints over 72 hours. 3D7 is a lab-adapted strain used in routine culture. Th.026.09 is a recent clinical isolate from Senegal. **(d)** RBC phenotypes were measured using complete blood counts with RBC indices, osmotic fragility tests, and ektacytometry.

### *P. falciparum* replication rates vary widely among non-carrier RBCs

To determine the variation in *P. falciparum* fitness among samples with different genotypes, we performed invasion and growth assays with two strains of parasite: a laboratory adapted strain (3D7) and a recent clinical isolate from Senegal (Th.026.09). We observed a wide range of *P. falciparum* growth rates among RBC samples, especially among non-carriers that lacked known disease alleles (Figure 2A-C). Each strain’s growth rate is defined here as parasite multiplication over a full 48-hour cycle in donor RBCs (Figure 1C), with the mean value for non-carriers set to 100% after normalization. Briefly, we used a repeated control RBC sample (Figure 2, gray circles) and other batch-specific factors to correct for variation in parasite growth across multiple experiments (see Methods). Among non-carriers, growth rates ranged from 64-136% for 3D7 (SD = 17.7%) and 76-128% for Th.026.09 (SD = 10.7%) (Figure 2A-B). Per-sample growth rates were strongly correlated between the two strains (Figure 2C, p < 3×10^-16^) and positively correlated when measured in different weeks (Figure S1A, *ρ*=0.34), demonstrating that these data capture real variation among donor RBCs. Furthermore, as expected (Friedman, 1978; Ifediba *et al*., 1985; Greene, 1993; Facer, 1995), we detected reductions in mean growth rate for both strains in RBCs carrying known disease alleles. These included severe G6PD deficiency (3D7, 60% of the non-carrier average, p<0.014; Th.026.09, 64%, p<0.014), α-thalassemia trait (3D7, 75%, p=0.027; Th.026.09, 81%, p = 0.077), HbAS (3D7, 16%, p=1.05×10^-7^; Th.026.09, 19%, p =1.2×10^-4^), and HE (3D7, 18%, p<0.014; Th.026.09, 33%, p<0.014). Notably, the wide distribution of growth rates for non-carrier RBCs had considerable overlap with the growth rates in carrier RBCs. Only the HbAS and HE samples fell entirely outside the non-carrier range. This observation implies the existence of previously unknown RBC variation that impacts *P. falciparum* growth, which may have effect sizes comparable to the milder known disease alleles.

**Figure 2.**
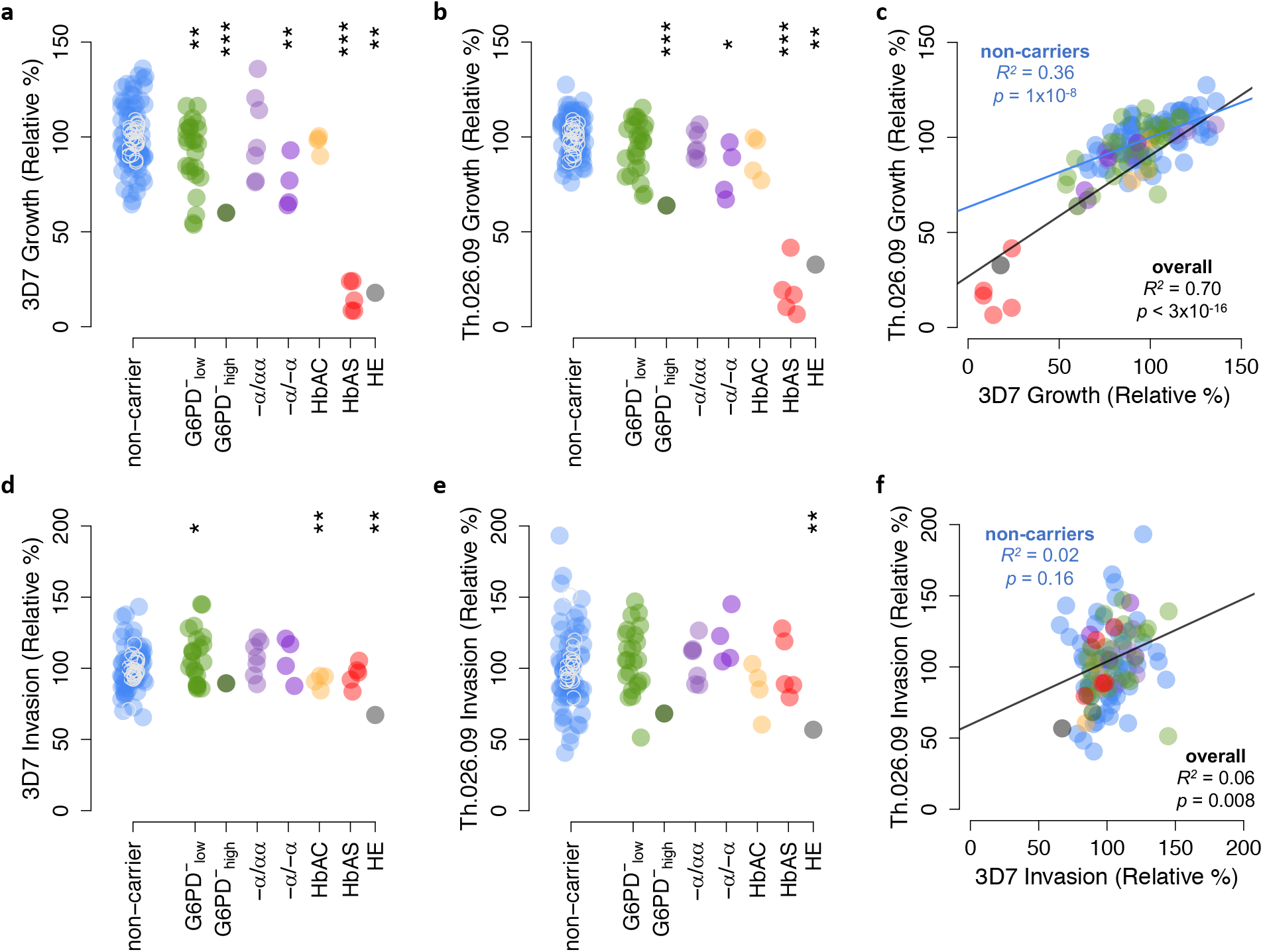
*P. falciparum* replication rate varies widely among donor RBCs. **(a-b)** Growth of *P. falciparum* strain 3D7 **(a)** or clinical isolate Th026.09 **(b)** over a full 48-hour cycle in donor RBCs_(hours 24-72 of assay). Growth is presented relative to the average non-carrier rate after correction for batch effects, including comparison to a repeated RBC control (Methods). Repeated measurements of the same RBC control over the course of the experiment are shown as open gray circles. **(c)** Correlation in per-sample growth rates between the two *P. falciparum* strains. **(d-e)** Invasion efficiency of *P. falciparum* strain 3D7 **(d)** or clinical isolate Th026.09 **(e)** after a 24-hour incubation with donor RBCs, normalized in the same way as growth. Repeated measurements of the RBC control over time are shown as gray circles. **(f)** Correlation in per-sample invasion success for the two *P. falciparum* strains. Colors in **(c)** and **(f)** correspond to the classifications in other panels; *R^2^* and *p*-values are derived from OLS regression. *p<0.1; **p<0.05; ***p<0.01.

We observed a similarly wide range in the efficiency of *P. falciparum* invasion into donor RBCs (Figure 2D-F). Invasion is defined here as the fold-change in parasitemia over the first 24 hours of the assay, when parasites previously maintained in standard culture conditions egressed and invaded new donor RBCs (Figure 1C). Among non-carriers, invasion rates ranged from 66-173% for 3D7 (SD=15.8%) and 41-193% for Th.026.09 (SD=28.9%) (Figure 2D-E). Compared to growth rates, no disease alleles conferred protection against invasion that was extreme enough to fall outside the broad non-carrier range. In 3D7 only, HbAC was associated with a 9% decrease in invasion (p=0.019) and mild G6PD deficiency was associated with a 7% increase (p=0.072). Only HE had a significant effect on the invasion efficiency of both strains (3D7, 67% of the non-carrier average, p<0.014; Th.026.09, 57%, p<0.014). There was little correlation between the invasion efficiencies of the two parasite strains in individual donor samples (Figure 2F), potentially reflecting strain-specific differences in the pathways used for invasion (Wright and Rayner, 2014). However, as we also observed greater variability between repeated samples for invasion than for growth (Figure S1), the invasion results may be influenced by greater experimental noise.

### RBC phenotypes vary widely among non-carriers

To assess phenotypic variation across donor RBCs, we measured 19 common indices of RBC size and hemoglobin content from complete blood counts using an Advia hematology analyzer (Figure 3A-D; Figure S2). Mean cellular volume (MCV) and hemoglobin mass (MCH) are closely related traits, which can be represented together as cellular hemoglobin concentration (CHCM) or the fraction of RBCs with ‘normal’ hemoglobin and volume indices (M5). As expected, each known disease allele was associated with a distinct set of RBC abnormalities. These included elevated CHCM for HbAC (p=0.033), consistent with dehydration, and very low MCV (p=8.7×10^-5^) and MCH (p=4.6×10^-7^) for α-thalassemia trait (αα/--), consistent with microcytic anemia (Galanello and Cao, 2011). RBCs from the HE patient also had very low MCV and MCH (both p<0.014), reflecting the membrane breakage and volume loss characteristic of this disease. However, for all of these phenotypic measures, we also observed broad distributions in non-carriers that overlapped the distributions of most carriers (Figure 3A-D; Figure S2). Notably, the breadth of the non-carrier distribution for each phenotype was large (e.g., 24 fL range for MCV) compared to the average difference between Africans and Caucasians (e.g., 3-5 fL(Beutler and West, 2005; Lo *et al*., 2011)). This wide diversity and substantial overlap between non-carrier and carrier traits suggest that ‘healthy’ RBCs exist on the same phenotypic continuum as RBCs carrying known disease alleles.

**Figure 3.**
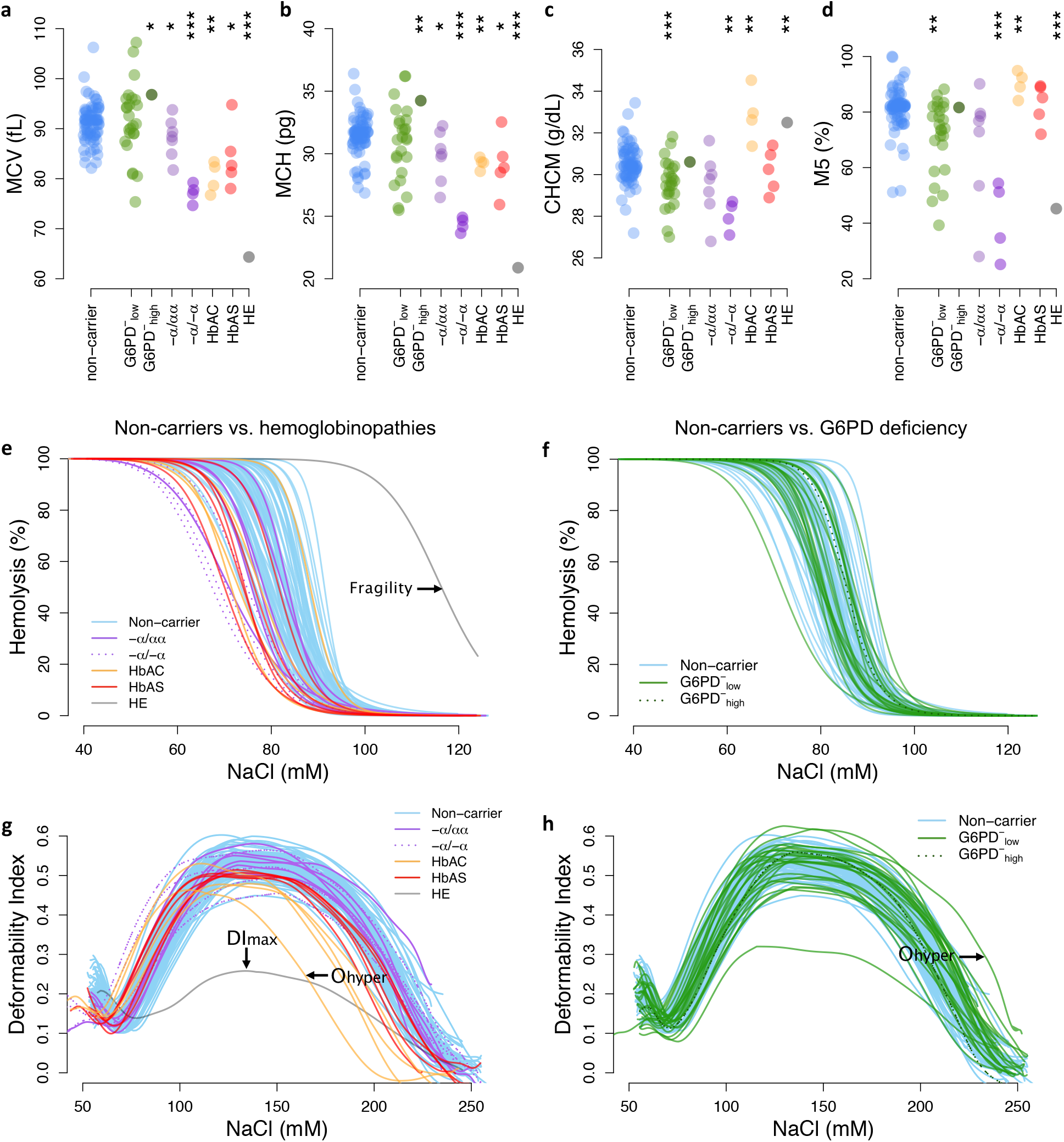
Red cell phenotypes that are abnormal in carriers also vary widely among non-carriers. **MCV:** mean RBC volume. **MCH:** mean cellular hemoglobin. **CHCM:** cellular hemoglobin concentration. **M5:** fraction of RBCs with ‘normal’ indices of hemoglobin concentration (28-41 g/dL) and volume (60-120 fL). **Fragility:** Tendency to lyse when osmotically overhydrated, defined as [NaCl] causing 50% lysis. **DI_max_:** Maximum membrane deformability under physiological salt conditions. **O_hyper_:** Tendency to resist osmotic dehydration and loss of deformability, defined as the hypertonic [NaCl] at which deformability is half maximum. *p<0.1; **p<0.05; ***p<0.01; tests and samples as in **Figure 2.**

We observed similar patterns of variation in RBC membrane fragility (Figure 3E-F) and membrane deformability (Figure 3G-H), as measured with osmotic fragility tests and osmotic gradient ektacytometry (see Methods). Both sets of curves represent RBC tolerance to osmotic stress, which can result in swelling and lysis (fragility, Figure 3E-F) or dehydration and decreased deformability (O_hyper_, Figure 3G-H). Specific hemoglobinopathies were associated with moderate to strong reductions in fragility (αα/α- p=0.039; αα/-- p=0.003; HbAS p=0.014), deformability (DI_max_; HbAC p=0.007; HbAS p=4.4×10^-6^), and/or resistance to loss of deformability when dehydrated (O_hyper_; HbAC p=0.019; HbAS p=0.041). HE cells were both extremely fragile (p<0.014) and extremely non-deformable (p<0.014). In non-carriers, the distributions for all membrane measures were wide, continuous, and overlapped the distribution of most carriers (Figure 3E-H; Figure S3). Overall, these data demonstrate that multiple phenotypic alterations associated with RBC disease alleles are also present in non-carrier RBCs.

### Non-carrier variation in RBC phenotypes predicts *P. falciparum* replication rate

To identify sets of phenotypes associated with *P. falciparum* replication in non-carrier RBCs, we used a machine learning method called Lasso (least absolute shrinkage and selection operator) that performs regularization and variable selection (Tibshirani and Tibshirani, 1994) (see Methods). Briefly, Lasso shrinks the regression coefficients for some possible predictors (in this case, phenotypes) to zero, in order to obtain a subset of predictors that minimizes prediction error. This method is well-suited for datasets in which possible predictors are correlated, as are standard measurements of RBC size, hemoglobin, and membrane dynamics. For each trait in each predictive subset selected by Lasso, we then applied univariate OLS regression to more accurately estimate the sign of its effect on all measured components of parasite fitness. The highest-confidence results are summarized in Figure 4A, with complete details provided in Table S2.

**Figure 4.**
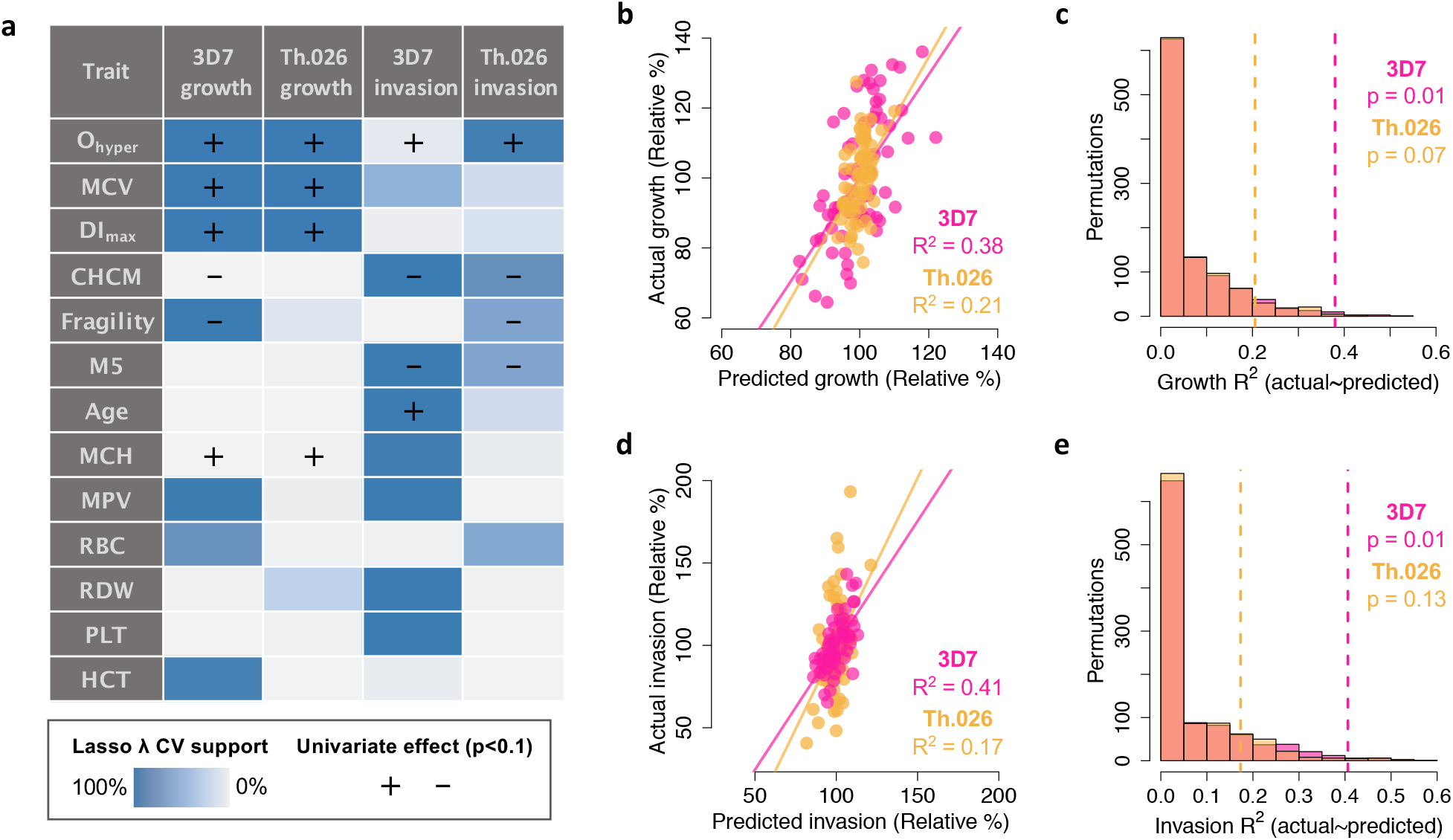
RBC phenotypes predict *P. falciparum* fitness in non-carriers. **(a)** Phenotypes shown were selected by Lasso in at least one of four models (columns) with at least 60% cross-validation (CV) support, as indicated by blue shading. **(+/-)** shows the direction of effect if the phenotype was significantly correlated (*p*<0.1) with the parasite fitness component in a separate, univariate linear model. **(b)** Lasso model predictions of parasite growth as a function of RBC phenotypes are plotted against measured growth values. Each point represents one non-carrier. The solid line shows a perfect 1:1 fit; *R^2^* indicates the proportion of variance explained by Lasso predictions. **(c)** For 1000 permutations of the growth data, the histograms show the proportion of variance explained by the Lasso modeling procedure. Dashed lines indicate *R^2^* for the actual data. *p*-values indicate the fraction of permutations that explain at least as much variance as the real data. **(d,e)** As in **(b,c)**, but for *P. falciparum* invasion. **O_hyper_:** Tendency to resist osmotic dehydration and loss of deformability, defined as the hypertonic [NaCl] at which deformability is half maximum (mM). **MCV:** mean RBC volume (fL). **DI_max_:** Maximum membrane deformability under physiological salt conditions. **CHCM:** cellular hemoglobin concentration (g/dL). **Fragility:** Tendency to lyse when osmotically overhydrated, defined as [NaCl] causing 50% lysis (mM). **M5:** fraction of RBCs with ‘normal’ indices of hemoglobin concentration (28-41 g/dL) and volume (60-120 fL) (%). **MCH:** mean corpuscular hemoglobin (pg/RBC). **MPV:** mean platelet volume (fL). **RBC**: red cell number (x10^6^/uL). **PLT:** platelet number (x10^3^/uL).). **HCT:** hematocrit (%).

*P. falciparum* fitness in non-carrier RBCs was strongly predicted by variation in several RBC traits related to deformability, dehydration, volume, and hemoglobin content. Among 24 tested phenotypes, the most important trait was the ektacytometry parameter O_hyper_, which represents a cell’s tendency to retain deformability in the face of dehydration (Figure 3G). In univariate models, non-carrier RBCs with the largest O_hyper_ values supported 1.2-1.5X faster parasite growth (3D7 1.47X, p=0.003; Th.026.09 1.24X, p=0.005) and 1.4-2.0X more effective invasion (3D7 1.36X, p=0.018; Th.026.09 1.94X, p=0.006) than RBCs with the smallest O_hyper_ values (Table S2; Figure S5A). Consistent with this result, *P. falciparum* replication was relatively inhibited in RBCs that were more dehydrated at baseline, as indicated by higher CHCM (3D7 invasion p=0.007; 3D7 growth p=0.042; Th.026.09 invasion p=0.006; Th.026.09 growth p=0.36). Parasites also grew faster in RBCs with larger mean volume, or MCV (3D7 p=0.001; Th.026.09, p=0.001); a greater mass of hemoglobin per cell, or MCH (3D7 p=0.073; Th.026.09 p=0.019); and more deformable membranes, measured by DI_max_ (3D7 p=0.0006; Th.026.09 p=0.006). Some measures of parasite fitness were also reduced in RBCs with more fragile membranes (3D7 invasion p=0.65; 3D7 growth p=0.003; Th.026.09 invasion p=0.067; Th.026.09 growth p=0.14). Additional phenotypes related to platelets and RBC density were selected for some models, but the direction of their effects was unclear since they were not significant when considered individually. These results indicate that common, non-pathogenic variation in RBC size, membrane dynamics, and other correlated traits have meaningful effects on *P. falciparum* replication rate in RBCs.

Together, 13 phenotype measurements explained approximately 40% of the variation in 3D7 growth and invasion in non-carriers (Figure 4B,D). To test for overfitting, we applied the same model-fitting procedure many times to randomly shuffled parasite data, which usually failed to produce predictive models (i.e., R^2^=0, Figure 4C,E). These permutation tests found that real data explained, on average, 6.5X the variance in 3D7 growth and 6.3X the variance in 3D7 invasion than expected by chance (both p=0.01, Figure 4C,E). For Th.026.09, the phenotypes selected by Lasso explained 17-21% of the total variation in parasite fitness (Figure 4B), which was marginally better than expected by chance for growth (3.7X, p=0.07, Figure 4C) and marginally worse than expected for invasion (3.0X, p=0.13, Figure 4E). The weaker result for Th.026.09 may be explained by its smaller dynamic growth range (Figure 2A-B), perhaps because clinical isolates are relatively poorly adapted to laboratory conditions. Overall, these results demonstrate that multiple, variable phenotypes impact *P. falciparum* susceptibility in healthy RBCs. Non-carrier cells that are less hospitable to parasites share specific traits with RBCs that carry disease alleles, including smaller size, decreased deformability, and an increased tendency to lose deformability when dehydrating.

### A small set of RBC genotypes predicts *P. falciparum* replication in non-carriers

Next, we tested whether non-carrier genotypes derived from exome sequencing could improve our predictions of *P. falciparum* replication rate. To limit false positives and increase statistical power (Ioannidis, 2005; Flynn, Hurvich and Simonoff, 2017), we focused on 23 candidate RBC proteins that have been previously associated with *P. falciparum* resistance in the literature (Table S3). These proteins contained 106 unlinked genetic variants (pairwise *r^2^* < 0.1) among the 73 non-carriers. Along with these polymorphisms and phenotypes described above, we also treated the top 10 principal components (PCs) of 1000 Genomes population structure as possible predictors of *P. falciparum* fitness.

The addition of genetic factors substantially improved all four models of parasite fitness. Across strains, 73-82% of the parasite variation in non-carriers was explained by a total of 47 genetic and phenotypic variables selected by Lasso (Figure 5A; Table S4). For 3D7 and Th.026.09 respectively, polymorphisms in 14 of 23 malaria-related genes explained 71% and 41% of the growth variation in non-carriers (Figure 5B). These models were significant by permutation (both p=0.011; Figure S6AB), indicating that the results were unlikely to be due to overfitting. Nonetheless, to confirm that the genetic signal was specific to our set of 23 malaria-related genes, we used polymorphisms from 1000 sets of random genes from the RBC proteome (Table S6) to build 1000 alternative models (see Methods). On average, variants from random RBC genes explained only 17.9% of the observed variation in *P. falciparum* growth for 3D7 (p=0.002) and 16.4% for Th.026.09 (p=0.047) (Figure 5B). In contrast to growth, the genotype/phenotype models of *P. falciparum* invasion (Figure 5C) were only marginally significant by permutation (3D7 p=0.09, Th.026.09 p=0.11, Figure S6C-D) and the genetic signal was not specific to the set of 23 malaria-related genes (3D7 p=0.26, Th.026.09 p=0.52, Figure 5D). Nonetheless, 4 of the 8 variants selected by Lasso for 3D7 invasion were also selected for 3D7 growth. Together, these results indicate that *P. falciparum* growth rate in RBCs is strongly determined by host genetic variation concentrated in a small number of genes.

**Figure 5.**
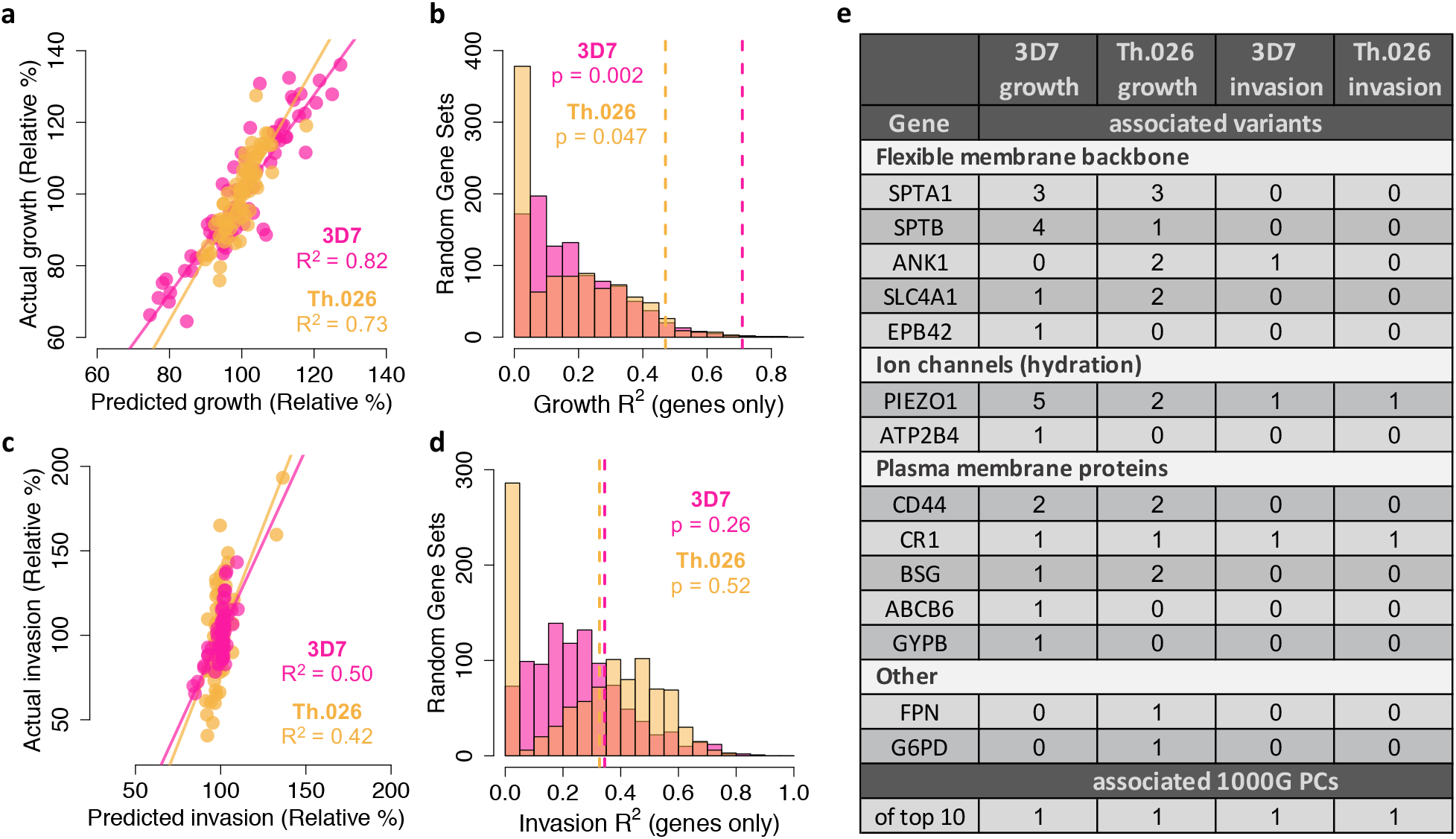
RBC genotypes substantially improve predictions of *P. falciparum* fitness in non-carriers. **(a)** Lasso model predictions of parasite growth as a function of RBC phenotypes, PCs of population structure, and variants in 23 malaria-related proteins (Table S3) are plotted against measured growth values. Each point represents one non-carrier. The solid line shows a perfect 1:1 fit; *R^2^* indicates the proportion of variance explained by Lasso-selected predictors. **(b)** For 1000 random sets of 23 genes drawn from the RBC proteome, the histograms show the variance explained *by variants within those genes* after applying the same Lasso modeling procedure as (a). Dashed lines indicate the variance explained by variants within the 23 malaria-related genes. *p*-values indicate the fraction of random gene sets that explain at least as much variance as the real data. **(c,d)** As in (a,b), but for *P. falciparum* invasion. **(e)** Variants and PCs counted in each cell were selected in the designated Lasso model with at least 70% cross-validation (CV) support. Several tested genes had 0 associated variants: *CD55, EPB41, HP, HBA1/2, GYPA, GYPC, GYPB*, and *HBB*. Details on lower-confidence variants are available in Table S4.

Of the top 31 variants associated by Lasso with *P. falciparum* growth, seventeen (55%) were synonymous (Table S4). This fraction did not significantly differ from the input set of 106 variants (p=0.64, 2-tailed z-score). Although 15/17 of these synonymous variants have no known, direct function, 7/15 are in linkage disequilibrium (*r^2^* > 0.1) with other variants that have been associated with RBC phenotypes by GWAS (Table S4). Similarly, 6 of the top 14 non-synonymous polymorphisms associated with *P. falciparum* growth have been previously linked to RBC or other traits (Table S4). This overlap with previous studies suggests that the variants associated with *P. falciparum* growth in this study are likely to tag functional genetic loci.

Nearly all of the top 31 growth-associated polymorphisms occurred in either (1) ion channel proteins, which regulate RBC hydration; (2) components of the flexible RBC membrane backbone; or (3) red cell plasma membrane proteins, including known invasion receptors (Figure 5E). In the first category, the highly polymorphic ion channel *PIEZO1* contained six polymorphisms associated with small (−3%) to moderate (−32%) reductions of *P. falciparum* growth rate (Table S4). The microsatellite variant *PIEZO1*-E756del, which has been a focus of several recent studies (Ilboudo *et al*., 2018; Ma *et al*., 2018; Rooks *et al*., 2019; Nguetse *et al*., 2020), predicted a moderate reduction in Th.026.09 growth (−8.2%, p=0.011) but was not related to RBC dehydration in these data (Table S5). We also detected one growth-associated variant in *ATP2B4*, which encodes the primary RBC calcium channel PMCA4b (−13% 3D7, p=0.006). This variant tags an *ATP2B4* haplotype implicated by GWAS in protection from severe malaria (Timmann *et al*., 2012; Lessard *et al*., 2017; Zámbó *et al*., 2017) and many RBC phenotypes (van der Harst *et al*., 2012; Li *et al*., 2013; Lessard *et al*., 2017; Lin, Brown and Machiela, 2020) including RDW, which was replicated in our data set (Table S4). *SPTA1* and *SPTB*, which encode the flexible spectrin backbone of RBCs, also contained several variants associated with the growth of at least one *P. falciparum* strain, as did the structural linker proteins ANK1, SLC4A1, and EPB42 (Figure 5E; Table S4). Finally, we identified one polymorphism in ABCB6, two in CD44, and two in basigin (BSG) that were strongly associated with *P. falciparum* growth (Figure 5E; Table S4). These plasma membrane proteins have previously been implicated in *P. falciparum* invasion by *in vitro* genetic deficiency studies (Crosnier *et al*., 2011; Egan *et al*., 2015, 2018). Notably, two of the natural polymorphisms identified here are synonymous quantitative trait loci (QTL) for CD44 splicing (rs35356320) and BSG expression (rs4682) (GTEx Consortium, 2017). We also detected one variant in the RBC iron exporter *FPN* and one in *G6PD* with small effects on Th.026.09 growth (2.4%, p=0.015 and 3.4%, p=0.010 respectively; Table S4). No associated variants were detected in the other nine tested genes (Table S2), including three hemoglobin proteins, three glycophorins, *CD55, EPB41*, and *HP*. Together, these data demonstrate that *P. falciparum* growth rate in non-carrier RBCs is strongly determined by dozens of host genetic variants, which together shape the phenotypic distribution of red cell susceptibility.

In addition to these specific genetic variants, PCs of population structure explained 22-26% of the variation in non-carrier invasion and 10-12% of the variation in growth. Interestingly, neither PC1 nor PC2—which distinguish Africans from other populations (Figure 1A)—were related to parasite fitness. Instead, this signal of correlated variation was partially driven by a six-member family with unique ancestry in our sample (Figure 1A, white borders) and extreme parasite fitness values (Figure S6). Although the present study is limited by sample size, the association between global genetic PCs and *P. falciparum* fitness suggests that additional functional variants remain to be discovered in many populations.

### African ancestry does not predict *P. falciparum* resistance in red cells

Based on evidence from balanced disease alleles like HbAS, it has been suggested that anti-malarial selection has shaped polygenic red cell phenotypes in African populations as a whole (Goheen *et al*., 2016; Kanias *et al*., 2017; Ma *et al*., 2018). We tested this hypothesis by examining the correlation between African ancestry and *P. falciparum* replication rate in RBCs from non-carriers (Figure 6A-D). We found no evidence that these traits were related, apart from an weakly positive relationship between African ancestry and the invasion rate of Th.026.09, a clinical Senegalese strain (p=0.01, R^2^=0.09, Figure 6D). To understand this result, we next examined how key RBC phenotypes identified in this study (Figure 4A) vary with African ancestry. We found that greater African ancestry predicts reduced osmotic fragility (p= 9.03 x 10^-6^), increased hypotonic deformability (DI_min_, p=0.001), lower cellular hemoglobin concentration (CHCM, reflecting reduced dehydration, p=0.007), and a reduced fraction of RBCs with normal volume and hemoglobin content (M5, p=0.033). All of these traits actually predict *greater* red cell susceptibility to *P. falciparum* (Figure 4A), although together they explain less than 9% of the non-carrier variation in parasite fitness. The remaining key phenotypes (Figure 4A) do not vary with African ancestry, which may explain why African ancestry itself is mostly unassociated with *P. falciparum* fitness in non-carrier RBCs (Figure 6A-D).

**Figure 6.**
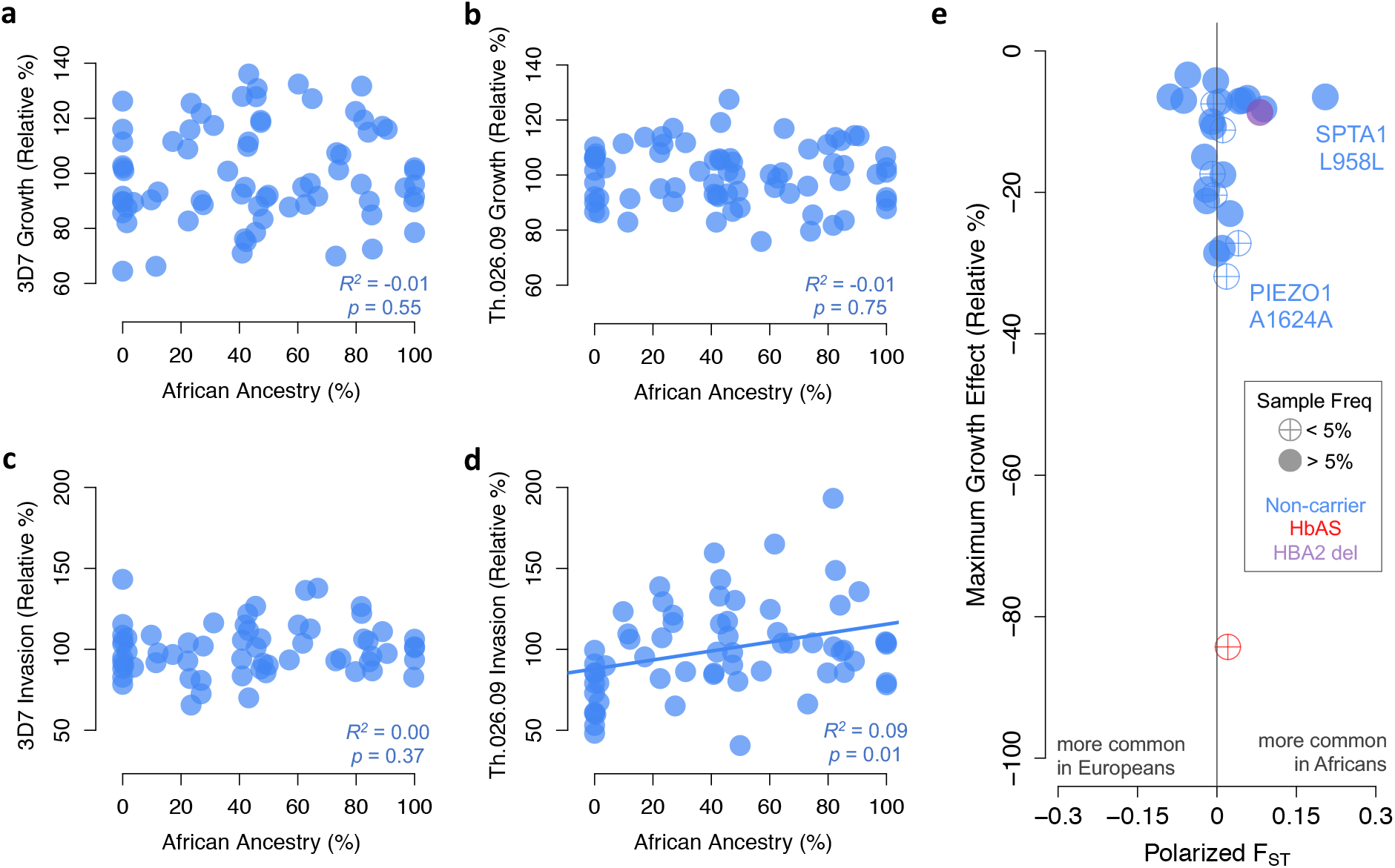
No evidence of widespread selection in Africa for slower *P. falciparum* replication or protective alleles in non-carriers. **(a-d)** Parasite growth and invasion measurements plotted against the exome-wide fraction of African ancestry, determined by comparison to 1000 Genomes reference populations. Adjusted-*R^2^* and *p*-values are shown for OLS regression. **(e)** Blue points represent non-carrier alleles selected by Lasso that also have significant effects on parasite growth (*p*<0.1) in univariate linear models (Table S4). The maximum estimated effect across two *P. falciparum* strains is shown. Red and purple points represent HbAS and the HBA2 deletion in an additive model, respectively. F_ST_ was calculated between pan-African and pan-European samples in gnomAD (Methods). Other disease alleles are not shown because they do not function as single loci (G6PD^-^, HE) or do not affect growth (HbAC).

Next, we used allele frequency data from the global gnomAD collection (Karczewski *et al*., 2020) to test whether polymorphisms associated with *P. falciparum* fitness in RBCs (Table S4) vary between African and European populations. Geographical differences in malaria selection are hypothesized to have increased the frequency of hundreds or thousands of undiscovered anti-malarial alleles in Africa (Mackinnon *et al*., 2005; Williams, 2006), as has been shown for several variants causing common RBC disorders (Kwiatkowski, 2005). Here, aside from well-known disease variants, we find no evidence that African populations as a whole are enriched for RBC polymorphisms that impair *P. falciparum* growth in at least one strain (Figure 6E). Of 25 alleles with detectable protective effects, 52% are in fact more common in Europeans than Africans (p=0.95, binomial test). Besides HbAS, the polymorphism with the largest effect is a synonymous variant in PIEZO1 (−31.9%, p=0.001), which is found at <4% frequency in both Africans and Europeans. A few other protective polymorphisms, including two in *PIEZO1* and two in *SPTB*, are nearly fixed in both populations. We do observe one F_ST_ outlier variant, a synonymous allele at SPTA1 L958, which is substantially more common in Africa (70% frequency) than in Europe (25%) and has a mild protective effect against *P. falciparum* growth (−6.5%, p=0.027, Figure 6E). Given its high frequency, as well as its direct association with a number of RBC phenotypes in previous GWAS (Table S4), this synonymous SPTA1 allele could be a promising candidate for investigating anti-malarial selection in Africa. However, the total set of natural RBC variants that limit *P. falciparum* growth *in vitro* do not appear to support the hypothesis of widespread divergent selection related to malaria between Africans and Europeans.

## DISCUSSION

Healthy red blood cells (RBCs) harbor extensive phenotypic and genetic variation, both within and between human populations. In this work, we demonstrate that this variation modulates a wide range of RBC susceptibility to *P. falciparum* parasites. Our findings add to a growing understanding of the genetic and phenotypic basis of RBC resistance to *P. falciparum*, especially for RBCs that lack population-specific disease alleles. In addition to suggesting new targets for future malaria interventions, these findings challenge assumptions about the role of malaria selection in shaping human RBC diversity.

Since exponential replication of *P. falciparum* is a significant driver of malaria disease progression (Bejon *et al*., 2007), the ample variation that we observed in this trait *in vitro* could be relevant for clinical outcomes in endemic regions. Growth inhibition from HbAS, for example, reduces the risk of death from malaria by reducing parasite density in the blood (Allison, 1954; Luzzatto, 2012). While HbAS has an extreme effect size, the 3-fold range of parasite replication rates we found among non-carrier RBCs shares substantial overlap with RBCs carrying other protective variants. Although the physiologically complex basis of severe malaria (Okwa, 2012) makes it difficult to estimate the precise contribution of RBC factors to severe malaria risk, the genotypes and phenotypes we have associated with *P. falciparum* fitness may be promising targets for future therapeutic interventions.

We have shown here that widespread, ‘normal’ variation in RBC traits like volume, hydration status, and maximum deformability is associated with *P. falciparum* fitness in non-carrier RBCs. Reassuringly, these phenotypes are present to a stronger degree in RBCs that do carry disease alleles (Clark, Mohandas and Shohet, 1983; Mockenhaupt *et al*., 2000; Pengon *et al*., 2018). They are also consistent with published reports of extreme RBC traits that limit parasite replication. For example, (Tiffert *et al*., 2005) used an experimental gradient of ion concentrations to show that *P. falciparum* growth is strongly reduced in chemically dehydrated RBCs. Here, we found that naturally occurring, mildly dehydrated RBCs also supported less efficient parasite growth and invasion. Similarly, the rare Dantu variant of the glycophorin A/B receptors has been associated with increased RBC membrane tension, decreased *P. falciparum* invasion efficiency, and decreased risk of severe malaria (Field *et al*., 1994; Leffler *et al*., 2017; Kariuki *et al*., 2018). Our data on RBC membrane deformability and fragility support the same link between more flexible RBC membranes and greater *P. falciparum* success. Moreover, they expand upon previous studies by demonstrating for the first time that common, healthy phenotypic variation in RBCs contributes meaningfully to *P. falciparum* fitness.

In our linear models of *P. falciparum* growth, phenotypic variation was strongly outperformed by genetic variation in a small number of RBC proteins. This result implies the existence of additional RBC phenotypes that we did not measure (or did not measure with sufficient accuracy), but which have a genetic basis in a small number of genes important to *P. falciparum*. Approximately half of the polymorphisms we identified are non-synonymous, and may therefore exert direct effects on phenotypes like RBC membrane structure or ion transport. The abundances of RBC gene transcripts and proteins are other clear candidates for future phenotype studies, given that the other half of associated polymorphisms were synonymous. A recent metanalysis showed that ‘silent’ and ‘coding’ SNPs are equally likely to be associated with human disease, and moreover have similar effect sizes (Chen *et al*., 2010). Although synonymous SNPs may sometimes be linked to coding variants, some have been shown to directly impact mRNA expression, protein translation speed, and protein folding by altering transcription levels, codon usage, and mRNA stability (Sauna and Kimchi-Sarfaty, 2011). Synonymous SNPs that impact splicing, like rs35356320 in CD44, may also impact protein structure. Other conceivable RBC phenotypes, such as the dynamics of membrane modification during *P. falciparum* development, may only become evident in more detailed time course experiments. The true number of RBC phenotypes important to *P. falciparum* may be effectively infinite (Kinsler, Geiler-Samerotte and Petrov, 2020), making it very useful in practice that static genetic variation is so strongly predictive of parasite growth.

Overall, just 31 polymorphisms in 14 RBC genes explained the majority of the variation in *P. falciparum* growth among non-carriers. One reason that our explanatory power is so high is our focus on *P. falciparum* replication in RBCs, which is a relatively simple component of a larger, complex disease. This focus allowed us to use controlled, *in vitro* experiments to detect genetic variants with small to moderate effects. Another important explanation is our use of Lasso variable selection on just 106 polymorphisms in genes with strong existing links to malaria. This approach obviated the need to meet an exome-wide significance threshold, while still allowing for the discovery of novel, putatively functional alleles in well-known disease genes. Furthermore, we confirmed that this set of genes was enriched for signal compared to random sets of genes drawn from the exome-wide background. It is important to note that genetic linkage complicates the identification of the exact functional polymorphisms in any population sample (Sohail *et al*., 2019), and we cannot rule out that some variants identified here are merely linked to the true functional variants. Indeed, about half of the variants we identified occur in linkage blocks containing other SNPs associated with RBC traits in GWAS studies. In any case, we present significant evidence that loci containing 14 RBC genes are strongly enriched for polymorphisms with significant impacts on *P. falciparum* growth.

One of the unique aspects of our study is the participation of individuals with a range of African ancestry, defined by similarity to donors from five 1000 Genomes reference populations. We found that African ancestry was associated with RBC phenotypes that *improved* parasite fitness, particularly for the clinical Senegalese strain Th.026.09. In the future, it would be very interesting to test for local parasite adaptation to human RBCs using *P. falciparum* strains and RBC samples from around the globe. We also found that the polymorphisms associated with *P. falciparum* growth by Lasso are not enriched in African populations included in the gnomAD database of human variation. This is not consistent with African-specific selection on these smaller-effect variants, despite the famous examples provided by RBC disease alleles. This unexpected result for ‘healthy’ variation has at least three possible explanations. First, relatively small-effect alleles may not have had sufficient time to increase in frequency since *P. falciparum* began expanding in humans some 5,000-10,000 years ago (Sundararaman *et al*., 2016; Otto *et al*., 2018). Second, the complexity of severe malaria could mean that the variants discovered here do not ultimately impact disease outcome, especially relative to known disease variants. Third, human adaptation may be too local to detect with coarse-grain sampling of Sub-Saharan African genetic diversity (e.g (Pankratov *et al*., 2020)). Our data do suggest, however, that few RBC alleles remain to be discovered that are both widespread in Africa and have large effects on *P. falciparum* proliferation in RBCs (e.g. (Ma *et al*., 2018)).

More broadly, these data show that it is inaccurate to make assumptions about RBC susceptibility to *P. falciparum* based on a person’s race or continental ancestry. These kinds of hypotheses (Williams, 2006; Goheen *et al*., 2016; Kanias *et al*., 2017; Ma *et al*., 2018) are based on well-known examples of balanced disease alleles, but our data suggest that these examples are exceptions rather than the rule. It is important to recognize that, at least in biology classes, the use of racially-based genetic examples increases the belief that genes encode absolute, functional differences between races (Donovan, 2017; Sparks, Baldwin and Darner, 2020). This perspective neglects the fact that >90% of genetic differences among humans occur within populations, rather than across them (Rosenberg, 2011). In this study, global population structure captured by PCs showed that non-African variation has its own distinct impacts on *P. falciparum* replication in RBCs. These data are an important reminder that most human genetic diversity is a result of demography, not population-specific selection (Coop *et al*., 2009; Hofer *et al*., 2009).

In conclusion, this study demonstrates that substantial phenotypic and genetic diversity in healthy human RBCs impacts the replication of malaria parasites. Whether or not this diversity is shaped by malaria selection, a better understanding of how *P. falciparum* biology is impacted by natural RBC variation can help lead to new therapies for one of humanity’s most important infectious diseases.

## ACKNOWLEDGMENTS

We gratefully acknowledge the invaluable participation of all volunteer blood donors. Nick Bondy, Sandra Larkin, Brian Fleischer, Ashley Dunn, Talal Seddik, Trung Pham, David Vu, and Spectrum Child Health provided crucial assistance in donor coordination and sample processing. *P. falciparum* strain Th.026.09 was kindly provided by Daouda Ndiaye and Sarah Volkman. For quantitative advice, we thank Grant Kinsler and the Stanford Statistics Consulting Group. This work was primarily supported by a Pilot Early Career award from the Stanford Maternal Child Health Research Institute and a Gabilan Faculty Award from the Stanford University School of Medicine Office of Faculty Development and Diversity. E.R.E. received additional support from the Stanford Center for Computational, Evolutionary, and Human Genomics. D.A.P. was funded through an NIH MIRA award 5R35GM118165-05. E.S.E. is a Tashia and John Morgridge Endowed Faculty Scholar in Pediatric Translational Medicine through the Stanford Maternal Child Health Research Institute. Local blood samples were drawn at the Stanford Clinical and Translational Research Unit, which is supported by CTSA Grant UL1 TR001085.

## AUTHOR CONTRIBUTIONS

Conceptualization, E.R.E., D.A.P., and E.S.E; Methodology, E.R.E., D.A.P., F.A.K., and E.S.E.; Investigation, E.R.E., F.A.K., C.L., and E.S.E., Formal Analysis, E.R.E.; Visualization, E.R.E., Writing-Original Draft, E.R.E.; Writing-Review & Editing, E.R.E., F.A.K., D.A.P., and E.S.E.; Funding Acquisition, E.R.E. and E.S.E.; Resources, F.A.K., D.A.P., and E.S.E; Supervision, D.A.P. and E.S.E.

## METHODS

### Sample collection and preparation

Subjects with no known history of RBC disorders were recruited to donate blood at the Stanford Clinical and Translational Research Unit. Written informed consent was obtained from each subject and/or their parent as part of a protocol approved by the Stanford University Institutional Review Board (#40479). To help control for weekly batch effects, Subject 1111 donated fresh blood for parasite assays each week. 11 other subjects donated blood on at least two different weeks. Whole blood samples from a hereditary elliptocytosis patient were obtained from Dr. Bertil Glader under a separate approved protocol (#14004) that permitted sample sharing among researchers. All samples were deidentified upon collection by labeling with a random four-digit number. Two samples were eventually removed from analysis based on a failed sequencing library (6449KD) and history of stem cell transplant (8715).

Whole blood was drawn into CPDA tubes and spun down within 36 hours to separate serum, buffy coat, and red blood cells. Red blood cells were washed and stored in RPMI-1640 medium (Sigma) supplemented with 25 mM HEPES, 50 mg/L hypoxanthine, and 2.42 mM sodium bicarbonate at 4°C. Buffy coat was transferred directly to cryotubes and stored at −80°C.

### Exome sequencing and genotype calling

Genomic DNA was isolated from frozen buffy coats using a DNeasy Blood and Tissue Kit (Qiagen). Libraries were prepared using a KAPA Hyperplus kit (Roche) and hybridized to human exome probes using the SeqCap EZ Prime Exome kit (Roche). The resulting exome libraries were sequenced with paired-end 150 bp Illumina reads on the HiSeq or NextSeq platforms at Admera Health (South Plainfield, NJ).

Reads were aligned to the hg38 human reference genome using bwa mem(Li, 2013), yielding an average coverage of 42X across targeted exome regions (excluding sample 6449KD). Variants were called using GATK best practices(Van der Auwera *et al*., 2013) and hard filtered with the following parameters: QD < 2.0, FS > 60.0, ReadPosRankSum < −2.5, SOR > 2.5, MQ < 55.0, MQRankSum < −1.0, DP < 500. To minimize the effects of sequencing errors, variants not present in 1000 Genomes, dbSNP_138, or the Mills indel collection(Mills *et al*., 2006) were discarded. Variants were also discarded if their frequencies in our sample did not fall between gnomAD African or European frequencies(Karczewski *et al*., 2020) and significantly differed from both groups’ frequencies by Fisher’s Exact Test (p≤0.05).

*PIEZO1* E756del was genotyped via PCR and Sanger sequencing according to a previously published protocol(Nguetse *et al*., 2020). To call deletion variants that cause α-thalassemia in the paralogous genes HBA2 and HBA1, we extracted reads from each .bam file that lacked any mismatches or soft-clipping and had MAPQ ≥ 13 (i.e., <5% chance of mapping error). Coverage with these well-mapped reads was calculated over the 73 and 81 bp of unique sequence in HBA2 and HBA1, respectively, and normalized to each sample’s exome-wide coverage. To determine which samples has unusually low coverage, we formed an *ad hoc* reference panel of seven donors who were unlikely to carry deletion alleles based on their normal MCH, MCV, and HGB and >96% exome-wide European ancestry(Weatherall, 2001). We called heterozygous HBA2 deletions when normalized coverage across three unique regions of the HBA2 gene was below the minimum reference value. Similarly, we called homozygous HBA2 deletions when normalized coverage across three unique regions of the HBA2 gene was less than half of the minimum reference value. This approach resulted in an estimated HBA2 copy number of 2.0 in the reference panel, 0.95 in eight putative heterozygotes and 0.12 in four putative homozygotes. The same method produced no evidence of HBA1 deletion in any sample.

### Variant classification and linkage pruning

Exonic variants in RefSeq genes were identified using ANNOVAR(Wang, Li and Hakonarson, 2010). Variants were classified into three categories: those within 23 malaria-related genes (Table S2); those within 887 other red blood cell proteins (Table S5) derived with a medium-confidence filter from the Red Blood Cell Collection database (rbcc.hegelab.org); and those within any other gene. Singleton variants were removed to avoid excessive false positives.

Linkage between all pairs of bi-allelic, exonic variants in our 121 genotyped samples was calculated using the --geno-r2 and --interchrom-geno-r2 functions in vcftools (Danecek *et al*., 2011). Variants in RBC genes that shared r^2^ > 0.1 with any variant in the 23-gene set were removed. Within the 23-gene set and RBC-gene set separately, non-carrier variants were ranked by the *p*-values of their OLS regression with all four parasite measures. Then, one variant was removed from each pair with r^2^ > 0.1, prioritizing retention in the following order: greater significance across models; non-synonymous protein change; higher frequency in our sample; and finally by random sampling. We report results from additive genetic models (genotypes coded 0/1/2), which performed as well or better than recessive (0/0/2) and dominant (0/2/2) models.

### Population analysis

The population ancestry of our donors was assessed by comparison with African, European, East Asian, and South Asian reference populations from the 1000 Genomes Project(The 1000 Genomes Project Consortium, 2015). Briefly, variants called from an hg38 alignment of the 1000 Genomes data (Lowy-Gallego *et al*., 2019) were filtered for concordance with the variants genotyped in this study. The --indep-pairwise command in PLINK(Purcell *et al*., 2007) was used to prune SNPs with r^2^ > 0.1 with any other SNP in a 50-SNP sliding window, producing 35,759 unlinked variants. These variants were analyzed in both PLINK --pca and in ADMIXTURE(Alexander and Lange, 2011) with K=4 for the 121 genotyped individuals in this study, alongside 2,458 individuals from 1000 Genomes. Pan-African and pan-European allele frequencies were obtained from gnomAD v3 (Karczewski *et al*., 2020). F_ST_ for specific alleles was calculated as (H_T_-H_S_)/H_T_ and then polarized, such that positive values indicate variants more common in Africa.

### *P. falciparum* culture and assays

*P. falciparum* strain 3D7 is a laboratory-adapted strain that was obtained from the Walter and Eliza Hall Institute (Melbourne, Australia) and routinely cultured in human erythrocytes obtained from the Stanford Blood Center. Th.026.09 is a clinical strain isolated from a patient in Senegal in 2009 that was then adapted to short-term culture (Park *et al*., 2012) and kindly provided by Daouda Ndiaye and Sarah Volkman. 3D7 was maintained at 2% hematocrit in RPMI-1640 supplemented with 25 mM HEPES, 50 mg/L hypoxanthine, 2.42 mM sodium bicarbonate and 4.31 mg/ml Albumax (Invitrogen) at 37°C in 5% CO2 and 1% O2. Th.026.09 was maintained in the same media, with the exception that half the Albumax was replaced with heat-inactivated human AB serum.

Parasite growth and invasion assays were performed using schizont-stage parasites isolated from routine culture using a MACS magnet (Miltenyi). Parasites were added at ~0.5% initial parasitemia to fresh erythrocytes suspended at 1% hematocrit in complete RPMI, as above. Parasites were cultured in each erythrocyte sample for three to five days in triplicate 100 uL wells. Parasitemia was determined on day 0, day 1 (24 hours), day 3 (72 hours), and in some cases day 5 (120 hours) by staining with SYBR Green 1 nucleic acid stain (Invitrogen, ThermoFisher Scientific, Eugene, OR, USA) at 1:2000 dilution in PBS/0.3% BSA for 20 minutes, followed by flow cytometry analysis on a MACSQuant flow cytometer (Miltenyi). Raw invasion rate was defined as the day 1 parasitemia divided by the day 0 parasitemia; raw growth rate was defined as the day 3 (or day 5) parasitemia divided by the day 1 (or day 3) parasitemia. Day 0 parasitemia was not measured in weeks 1-3, so invasion rate estimates are absent for these samples. The parasite assays failed for both strains in week 9 and for Th.026.09 in week 10, and so were repeated in weeks 10 and 11 with RBCs that had been stored for 1 or 2 weeks.

To correct for batch effects, such as week-to-week variation in *P. falciparum* replication, we extracted the residuals from a linear regression of the raw parasite values against up to four significantly related batch variables: (1) the raw values for control donor 1111 each week; (2) the parasitemia measured at the previous time point; (3) the age in weeks of the RBCs being measured; and (4) the experimenter performing the assays. Notably, there was no additional effect of ‘Week’ or the length of the experiment (i.e., 3 or 5 days) once the previous variables were regressed out. To convert these residuals (mean 0) to relative percents, we first transformed them with a linear model parameterized by data from control donor 1111 and the most extreme donors (HbAS for growth; G6PD-_high_ and HE for invasion), normalized by the 1111 rates from the appropriate week. These values were finally arithmetically adjusted so that the mean invasion and growth values for non-carriers was 100%.

### Red cell phenotyping and normalization

Complete blood count (CBC) data for RBCs, reticulocytes, and platelets were obtained with an Advia 120 hematology analyzer (Siemens, Laguna Hills, CA) at the Red Cell Laboratory at Children’s Hospital Oakland Research Institute. These data were: RBC, HGB, HCT, MCV, MCH, MCHC, CHCM, RDW, HDW, PLT, MPV, Reticulocyte number and percentage, and the fraction of RBCs in each of nine cells of the RBC matrix (M1-M9, with MCH cutoffs of 28 and 41 and MCV cutoffs of 60 and 120). Systematic biases were evident for some measures in certain weeks, but data from control donor 1111 were not available for all weeks. Therefore, CBC data were normalized such that the median value for non-carrier samples was equal across weeks.

Osmotic fragility tests were performed by incubating 20 μL of washed erythrocytes for 5 minutes in 500 μL solutions of NaCl in 14 concentrations: 7.17, 6.14, 5.73, 5.32, 4.91, 4.50, 4.30, 4.09, 3.89, 3.68, 3.27, 3.07, 2.66, and 2.46 g/L. Tubes were spun for 5 minutes at 1000g and 100 μL of supernatant was transferred to a 96-well plate in duplicate. Hemoglobin concentration was determined by adding 100 μL of Drabkin’s reagent (Ricca Chemical) to each well and measuring absorbance at OD_540nm_ with a Synergy H1 plate reader (Biotek). Relative lysis was determined by normalizing to the maximum hemoglobin concentration in the 14-tube series for each sample. After outlier points were manually removed, sigmoidal osmotic fragility curves were estimated under a self-starting logistic model in the nls package in R. Curves were summarized by the relative tonicity at which 50% lysis occurred and normalized within weekly batches, such that this value was equal for control sample 1111 across weeks.

Osmotic gradient ektacytometry (Clark, Mohandas and Shohet, 1983; Kuypers *et al*., 1990) was performed at the Red Cell Laboratory at Children’s Hospital Oakland Research Institute (CHORI). Red cell deformability estimates across a range of NaCl concentrations were fitted to a 20-parameter polynomial model to generate a smooth curve, which was manually verified to closely fit the data. Each curve was summarized with three standard points (Figure S3A)(Clark, Mohandas and Shohet, 1983), which were normalized such that the median x- and y-values of the three points was equal for non-carrier samples across weeks.

### Statistical analysis

Student’s t-test was used to compare trait values between non-carriers and each other group. Where n=1 (i.e., for G6PD_-high_ and HE), *p*-values were defined as the percentile of the non-carrier distribution. Significance in these tests was defined as *p*<0.1. OLS linear regression was performed with the lm function in R for all comparisons of two continuous variables, unless otherwise specified. Adjusted *R^2^* values are reported.

Lasso regression (Tibshirani and Tibshirani, 1994; Chatterjee, 2013) was performed with the cv.glmnet function in R. To account for moderate uncertainty in the choice of lambda (due to random division of data for cross-validations), each model was run 1000 times. For each predictor, we report the fraction of runs in which it was selected (e.g Figure 4A; Table S1; Table S3). The *R^2^* values reported between actual and predicted parasite measures are medians from 1000 runs. We noted that effect size estimates for each predictor varied considerably across runs; therefore, we used univariate OLS regression to separately estimate the effect size of each predictor selected at least once in at least one model.

The significance of each Lasso result was assessed by permutation. Parasite values were randomly shuffled 1000 times, preserving correlations among potential predictors. The median models for 6 cross-validations of each of the 1,000 permuted data sets were compared to the median model from the real data. To compare the 23 malaria-related genes to other sets of genes, a gene-specific *R^2^* for the target set was calculated using OLS with only specific variants (no phenotypes or PCs) selected in at least 70% of cross-validations. 1000 random sets of 23 genes were then generated from the RBC proteome (Table S5) and analyzed identically. *P*-values were calculated as the percentile of this permuted distribution in which the malaria-related gene set fell.

## LEAD CONTACT

Further information and requests for resources and reagents should be directed to and will be fulfilled by the Lead Contact, Elizabeth Egan.

## MATERIALS AVAILABILITY

This study did not generate any new unique reagents.

## DATA and CODE AVAILABILITY

The human sequence datasets generated during this study will be available at dbGAP (in progress). Phenotype data and code generated during this study is available at https://github.com/emily-ebel/RBC.

