## Supplementary material for "Common host variation drives malaria parasite fitness in healthy human red cells": SI: SI.docx (from another version of this preprint: version 2, 2020-10-21)

**Table S1. Linear models of batch effects on parasite fitness.**

**Table S2. All phenotypic predictors with non-zero Lasso support.** Phenotypes are ordered by their average Lasso cross-validation support across the four models.

**Table S3. Twenty-three RBC genes with strong links to malaria.**

**Table S4. All genetic and phenotypic predictors with non-zero Lasso support in a combined model.** Only predictors with >60% cross-validation support are shown in Figure 4e and Figure 5a.

**Table S5. Lack of association between RBC dehydration phenotypes and PIEZO1 rs59446030 or ATP2B4 rs1419114.**

**Table S6. List of proteins present in mature RBCs.** This list was derived from the Red Blood Cell Collection database (rbcc.hegelab.org) using a medium-confidence filter.

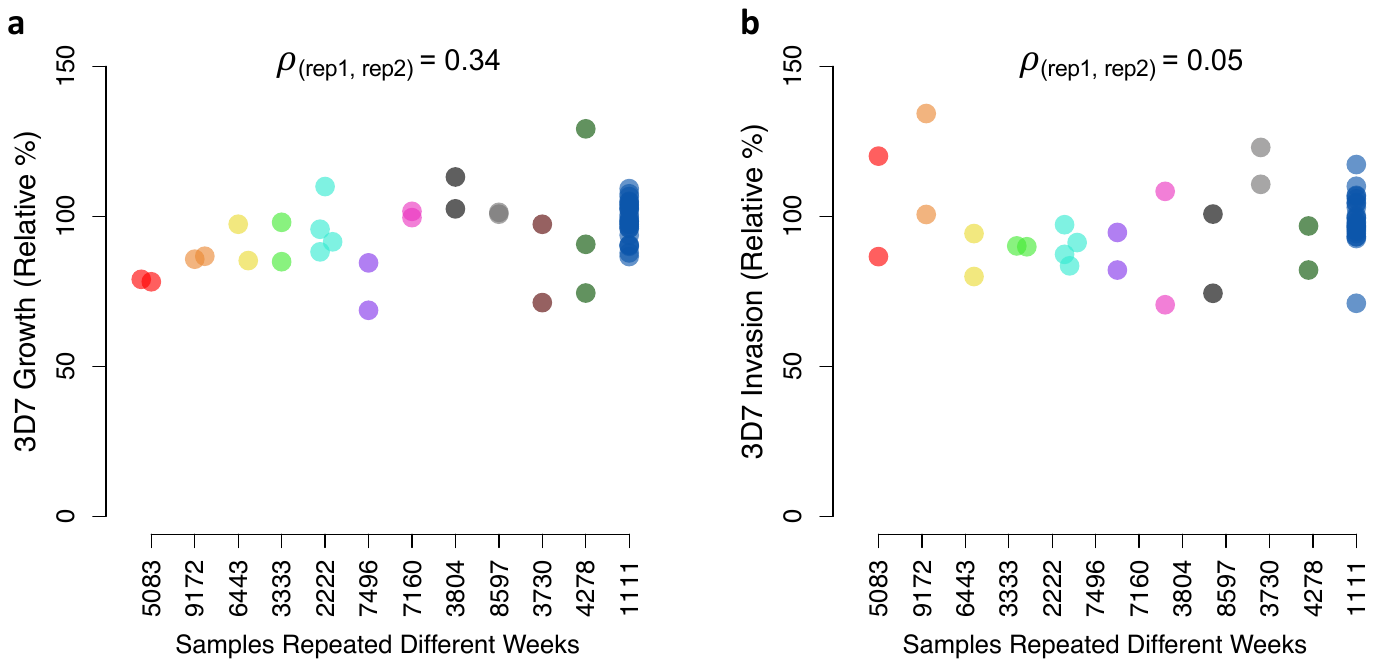

**Figure S1. Repeatability of parasite assays in the same donors over time.** 12 participants (including the weekly control, 1111) donated blood in multiple weeks for independent experiments. Pearson’s rho is calculated between the first and second assays and can range from -1 to 1. Growth and invasion are standardized as described in Methods. All repeated donors were non-carriers except 6443, who carried the HbAC allele; and 7160, 8597, and 4278, who carried alleles for mild G6PD deficiency.

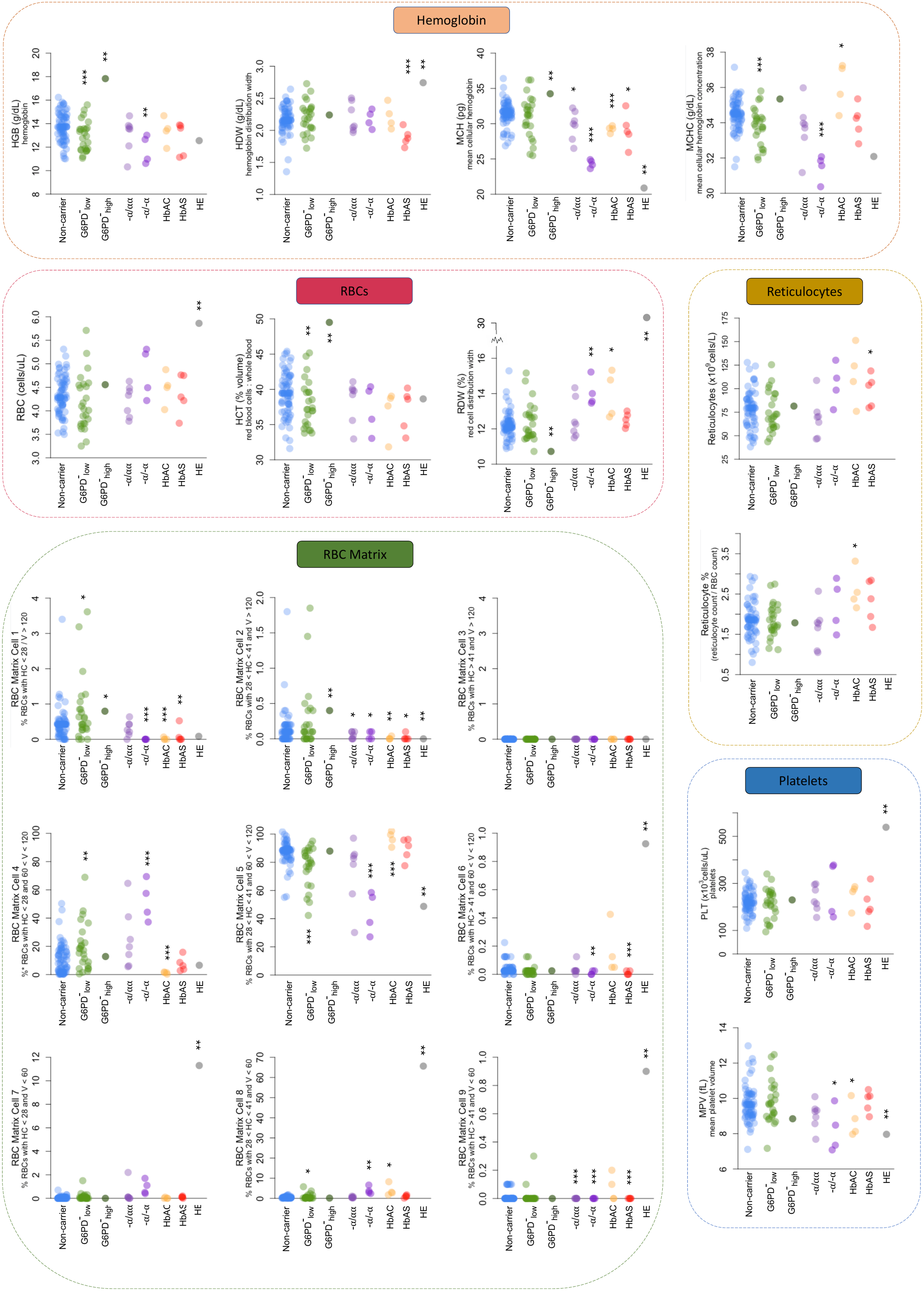
**Figure S2. Complete blood count data in donor RBCs.** Donor classifications and significance values as in Figures 1 and 2.

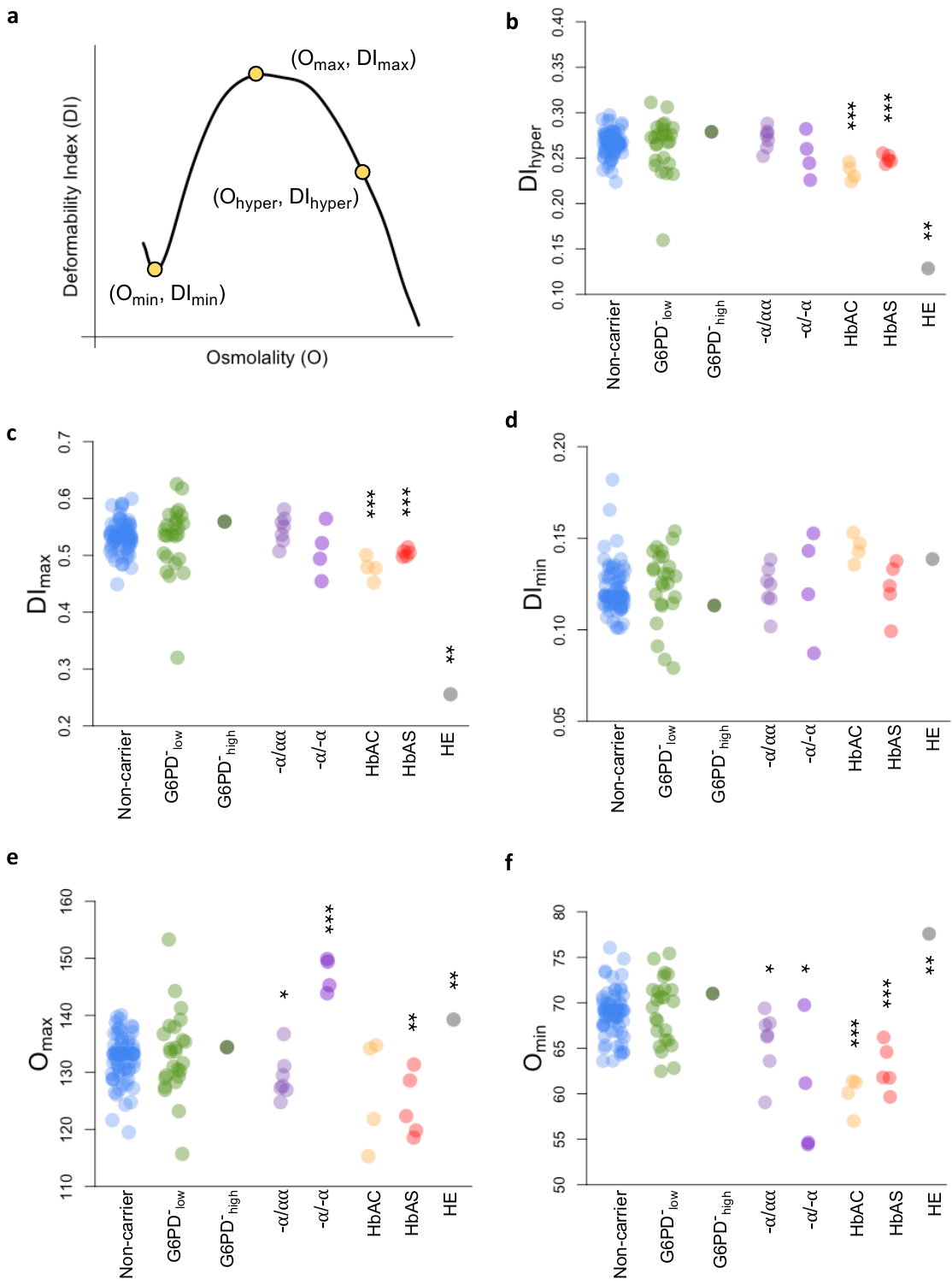

**Figure S3. Ektacytometry summary statistics.** Donor classifications and significance values as in Figures 1 and 2.

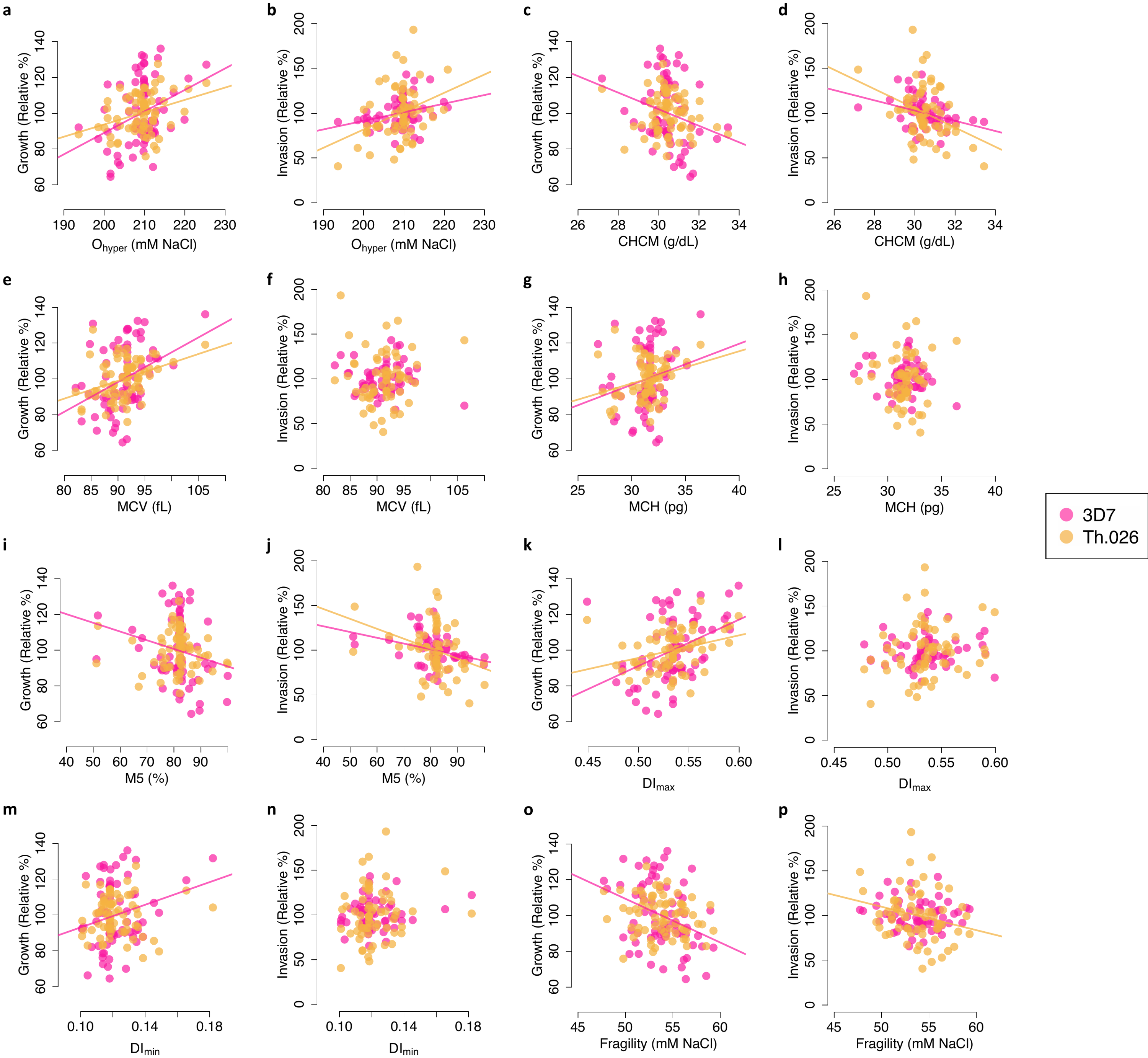

**Figure S4. Red cell phenotypes associated with *P. falciparum* replication in non-carriers.** Phenotypes are shown if they were selected by Lasso *and* are univariately related to at least one measure of parasite fitness. Lines of best fit are shown where p<0.1.

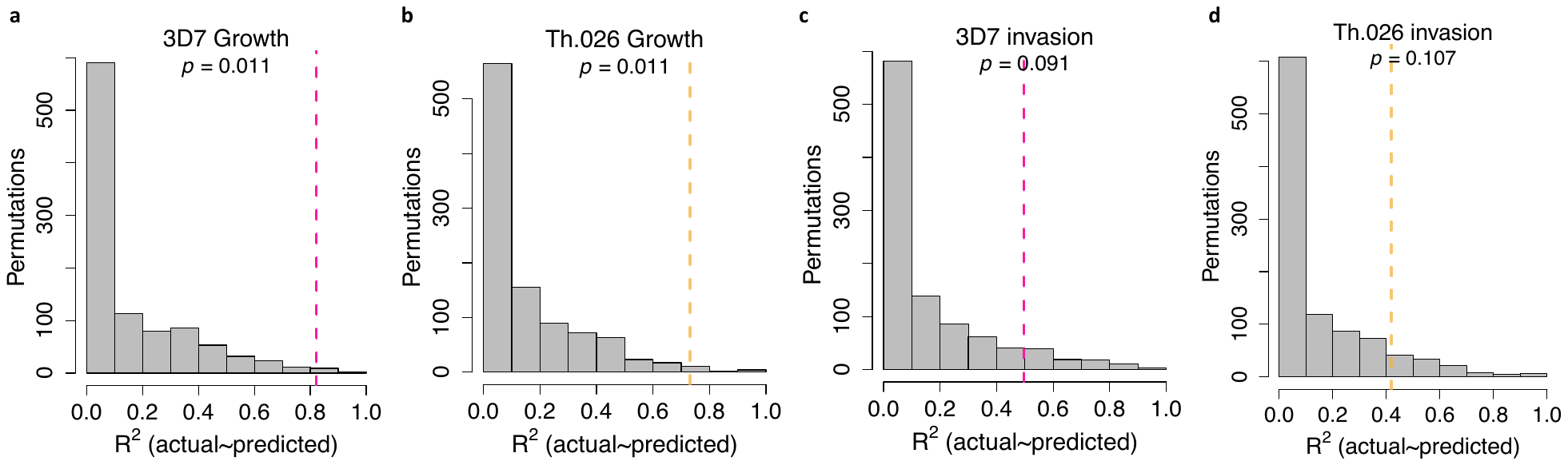

**Figure S5. Permutations testing for overfitting of Lasso models with genotype and phenotype data.** Dashed lines indicate the median *R^2^* for models built with real data. The histograms show the distribution of median *R^2^* for 1000 models built with permuted parasite data. *P*-values indicate the percentile of the permuted distribution in which the real data fall.

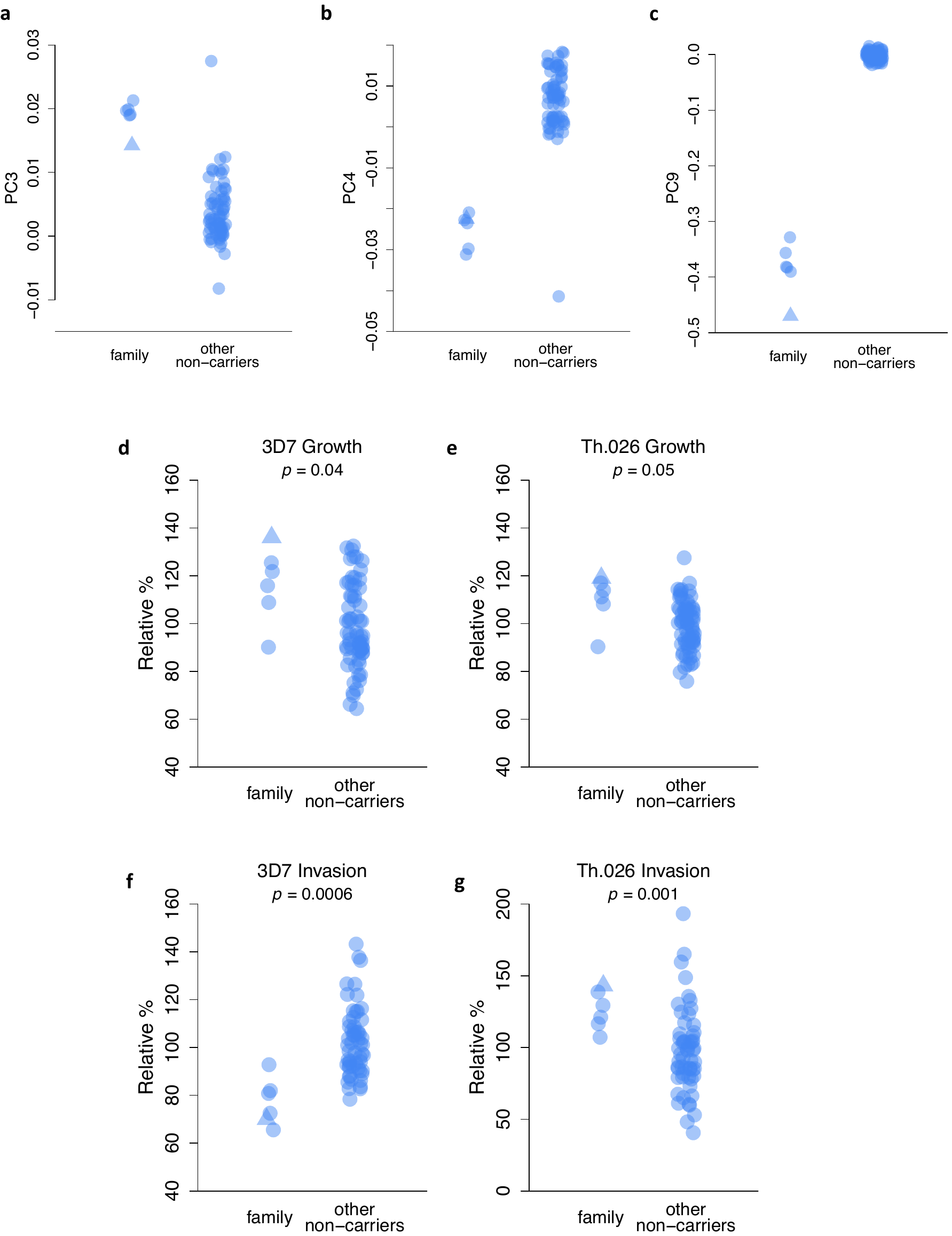

**Figure S6. A mother and her five children have unique ancestry and parasite susceptibilities compared to other non-carrier donors.** Data from the mother are indicated by a triangle. **(a-c)** Only PCs that distinguish the family from other non-carriers are shown. See also Fig 1. **(d-g)** *p*-values are derived from *t*-tests.

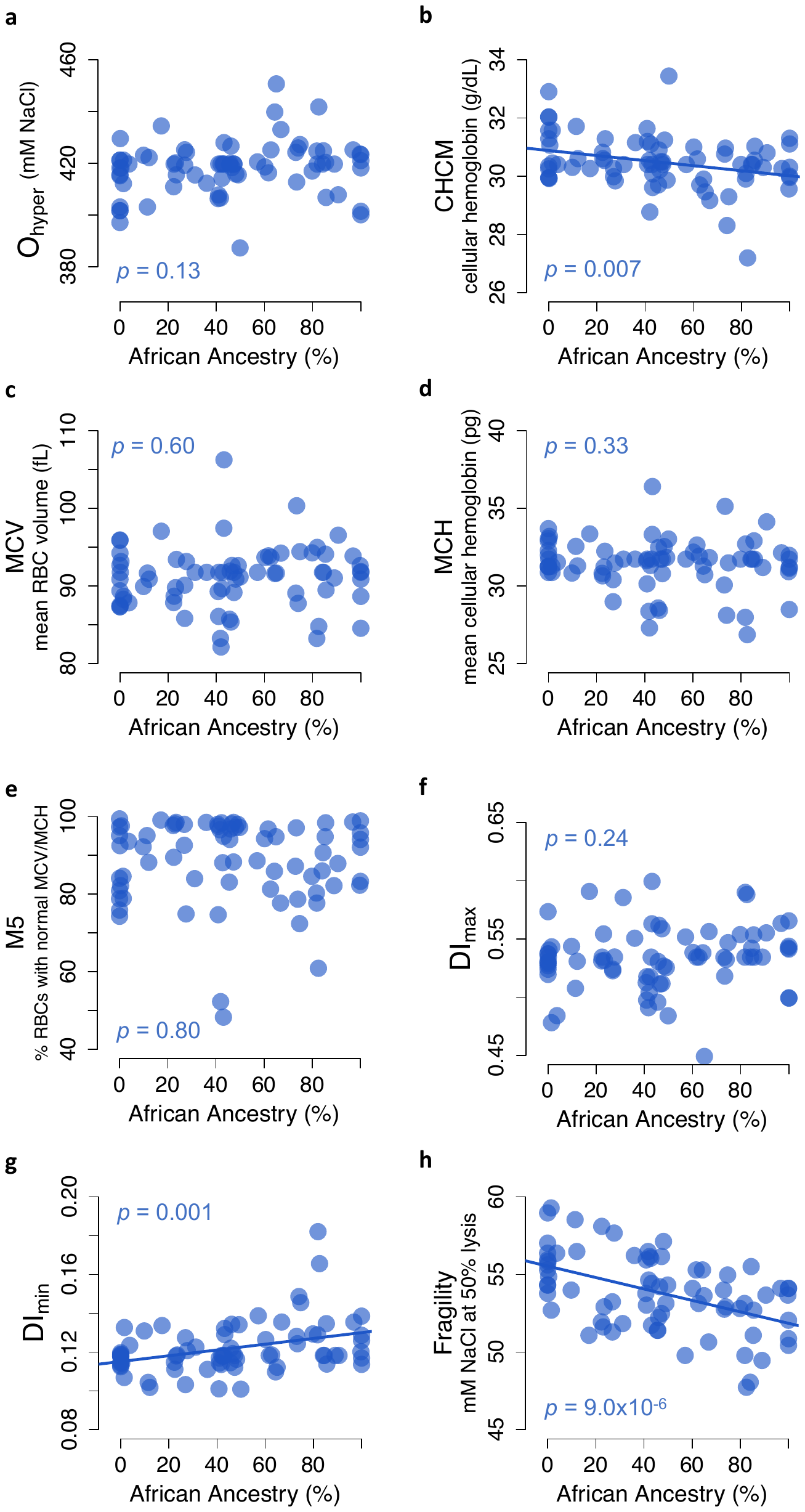

**Figure S7. RBC phenotypic variation versus exome-wide African ancestry (PC1).** *p*-values are from OLS regression. Lines of best fit are shown for associations where p<0.1.
